# Understanding West Nile virus spread: Mathematical modelling to determine the mechanisms with the most influence on infection dynamics

**DOI:** 10.1101/2022.02.15.480483

**Authors:** Elisa Fesce, Giovanni Marini, Roberto Rosà, Davide Lelli, Monica Pierangela Cerioli, Mario Chiari, Marco Farioli, Nicola Ferrari

**Author notes:** Department of Veterinary Medicine, Cambridge University, Madingley Road, CB3 0ES, Cambridge (UK). corresponding author, (EF).

## Abstract

West Nile disease is a vector-borne disease caused by West Nile virus (WNV), involving mosquitoes as vectors and birds as maintenance hosts. Humans and other mammals can be infected via mosquito bites, developing symptoms ranging from mild fever to severe neurological infection. Due to the worldwide spread of WNV, human infection risk is high in several countries. Nevertheless, there are still several knowledge gaps regarding WNV dynamics. Several aspects of transmission taking place between birds and mosquitoes, such as the length of the infectious period in birds or mosquito biting rates, are still not fully understood, and precise quantitative estimates are still lacking for the European species involved. This lack of knowledge affects the precision of parameter values when modelling the infection, consequently resulting in a potential impairment of the reliability of model simulations and predictions. Further investigations are thus needed to better understand these aspects, but field studies, especially those involving several wild species, such as in the case of WNV, can be challenging. Thus, it becomes crucial to identify which mechanisms most influence the dynamics of WNV transmission. In the present work, we propose a sensitivity analysis to identify the parameters that have the largest impact on the spread of the infection. Based on a mathematical model simulating WNV spread into the Lombardy region (northern Italy), the basic reproduction number of the infection was estimated and used to quantify infection spread into mosquitoes and birds. Then, we quantified how variations in the duration of the infectious period in birds, the mosquito biting rate on birds, and the competence and susceptibility to infection of different bird species might affect WNV transmission. Our study highlights that all considered parameters can affect the spread of WNV, but the duration of the infectious period in birds and mosquito biting rate are the most impactful, highlighting the need for further studies to better estimate these parameters. In addition, our study suggests that a WNV outbreak is very likely to occur in all areas with suitable temperatures, highlighting the wide area where WNV represents a serious risk for public health.

**Author Summary:** Infectious communicable diseases are currently one of the main burdens for human beings and public health. The comprehension of their spread and maintenance is one of the main goals to facilitate their control and eradication, but due to the complexity of their cycles and transmission mechanisms, obtaining this information is often difficult and demanding. The control of vector-borne diseases in particular represents an important and very complex challenge for public health. Mathematical models are suitable tools to investigate disease dynamics and their transmission mechanisms. To build a suitable model that can simulate transmission dynamics, a reliable and precise estimate of parameters for measuring transmission mechanisms is fundamental. We thus propose a sensitivity analysis of four unknown epidemiological parameters (bird recovery rate, mosquito biting rate, avian susceptibility to infection and avian competence to infection) that play a crucial role in driving West Nile virus (WNV) infection to determine which of them have the greatest impact on infection spread. This analysis allows us to identify the investigated parameters that need the most accurate estimate and to further investigate mechanisms that majorly affect WNV spread.

## Introduction

West Nile disease (WND) is an emergent vector-borne disease caused by West Nile virus (WNV), a single-stranded virus belonging to the Flavivirus genus [1]. Its cycle involves mosquitoes, mostly of the genus *Culex*, as vectors and diverse bird species as maintenance vertebrate hosts [2–5]. Although acting as dead-end hosts, several mammalian species, including humans, can be infected via mosquito bites and can develop symptoms ranging from mild fever to severe neurological disease [6,7]. Despite the low frequency of development of a severe illness (25% of infected persons develop symptoms [6]), recovery might take several weeks or months, and some effects on the central nervous system might be permanent [5,8]. In addition, recent diffusion of WNV in several countries in Europe and North America makes it one of the most widespread flaviviruses in the world [9]. In the Italian scenario in particular, WNV was detected for the first time in horses in the Tuscany region (central Italy) but is currently considered endemic in several Italian regions, making it a relevant public health issue. As a consequence, since 2013, health authorities have implemented an integrated surveillance system for early detection of its circulation. In particular, the increased number of cases registered in 2018 increased attention to the need to develop an adequate surveillance system to monitor WNV circulation, to comprehend mechanisms driving the spread of the infection and to fully understand the encountered spatiotemporal variability.

Despite that, the variety of bird species developing viraemia titres sufficient to infect feeding mosquitoes [10–12] and the differences in susceptibility and competence shown between and within families of birds [13–16] make WNV spread and diffusion still poorly understood, especially on the European continent [17]. Few studies have provided a complete analysis of different bird species’ viraemic responses, and the variety of the composition of local avian communities combined with the circulation of different WNV strains makes the extension of results to areas other than those tested inaccurate [13,15,16,18,19].

To investigate the spreading mechanisms of WNV, several mathematical modelling efforts have been made (e.g., [20–25]), but the abovementioned uncertainties might hinder the precision of parameter estimates, thus impairing the reliability of simulations obtained and the likelihood of simulated mechanisms. For this reason, the development of specific studies to investigate the epidemiological effects of different parameters is fundamental to improve the reliability of model simulations and our comprehension of WNV spread, especially in Europe, where information about species involved in WNV spread and their epidemiological characteristics are currently lacking [17,18,26]. When a disease involves several species, particularly wildlife species, as in the case of WNV, field studies can be difficult and demanding. Each parameter may affect WNV dynamics in a different way; in fact, small changes in some parameters can produce a tremendous variation in transmission dynamics, while larger changes in other parameters can scarcely affect it. For this reason, a precise quantification of those parameters having the largest effect on disease dynamics can aid the prioritization of future research. Our work aims to investigate the effects on infection spread of different estimates of four epidemiological parameters related to bird-mosquito-virus interactions whose values are relatively unknown. We pursued this aim by deploying a previously published computational framework [25] informed with entomological, climatic, and ornithological data gathered in the Lombardy region (northern Italy) between 2016 and 2018, and the investigated parameters were i) mosquito biting rate on birds, ii) avian competence, iii) avian recovery and iv) avian susceptibility to infection.

## Materials and methods

### Dataset and reference system

Entomological data were collected between 2016 and 2018 in the Lombardy region in northern Italy (see Fig 1). Mosquito abundance records and their WNV infection prevalence estimated by the Regional WNV mosquito surveillance program performed by Regione Lombardia and Istituto Zooprofilattico Sperimentale della Lombardia e dell’Emilia Romagna (IZSLER). Mosquito abundance and positivity for WNV were determined by field collection and PCR analyses [27]. According to national and regional guidelines for the entomological surveillance of WNV [27–30], field collection of mosquitoes was carried out every two weeks between May and September by the use of CO_2_ traps for capturing mosquitoes, covering an area of at most 400 km^2^ each (Fig 1). Daily mean temperature data for each quadrat, collected with ground stations, were obtained from ARPA Lombardia (Agenzia Regionale per la Protezione dell’Ambiente della Lombardia). We then divided the Lombardy region into 3 separate subregions, a northern subregion, a western subregion and an eastern subregion (red, green and blue coloured regions in Fig 1), accounting for differences in temperature, precipitation, geography, mosquito abundance and WNV circulation. Due to the absence of WND in the northern subregion, we investigated WNV transmission only through the eastern and western subregions. To estimate the number of birds composing the whole avian community at the beginning of the summer and, in particular, the number of magpies, known to be a susceptible and competent avian species [18,31], we considered the records of the avifauna census provided by Regione Lombardia [32]. Specifically, the number of magpies was estimated as a proportion of the total number of birds estimated by a kriging method calculating the number of individuals circulating in an area *A* = *π*\**r*^2^, where r is the average *Culex pipiens* flight range that was set to 500 m [23].

**Fig 1.**
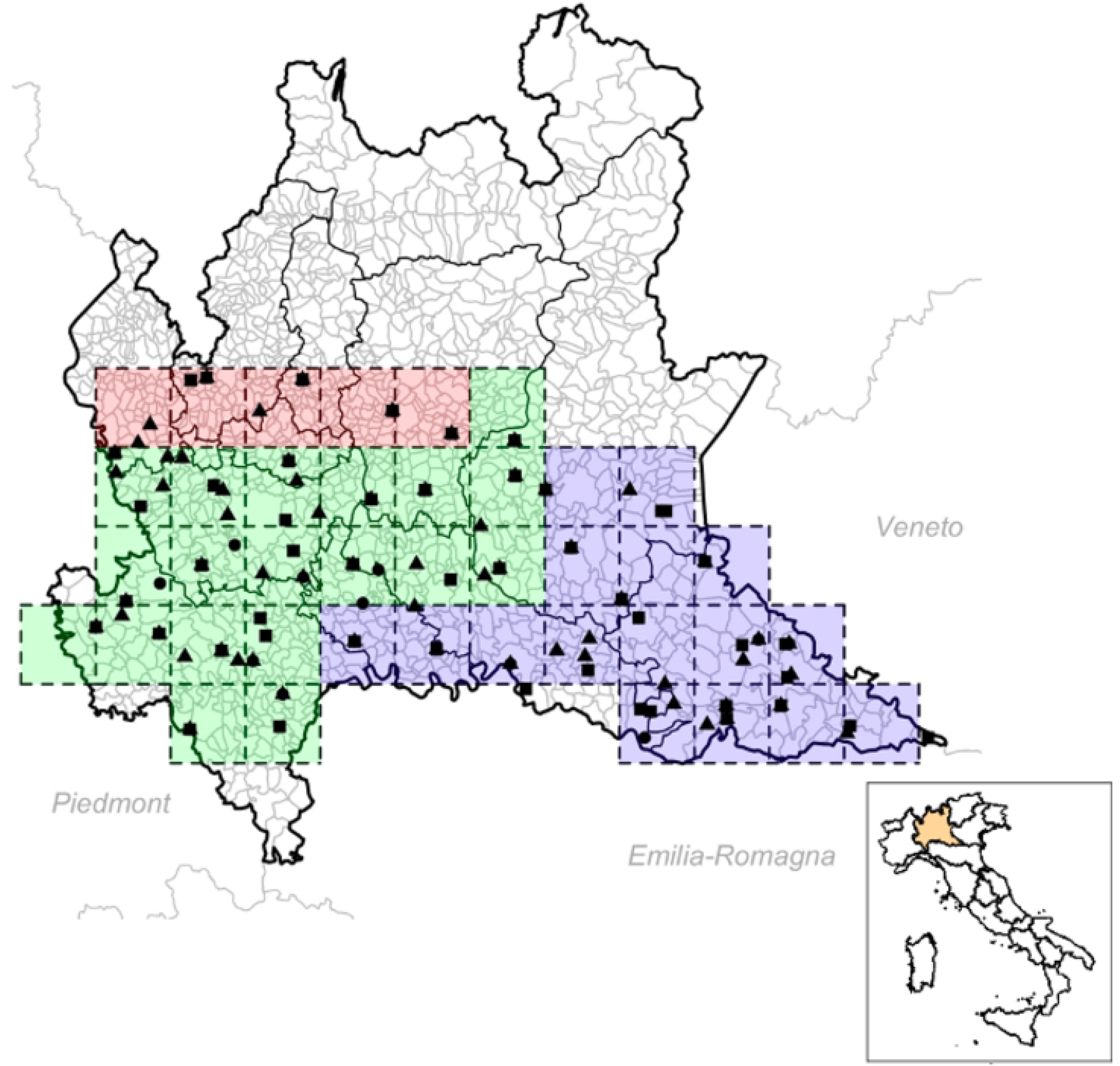
Capture site division proposed for the Lombardy region. Red quadrats represent the northern subregion, green quadrats represent the western subregion, and blue quadrats represent the eastern subregion. Black squares, circles and triangles represent mosquito capture sites for 2016, 2017 and 2018, respectively.

### Model structure

The modelling framework followed that proposed in Marini et al. [25] to investigate WNV dynamics in the Emilia Romagna region. First, the mosquito population dynamics were simulated through an *“entomological model”*, which provides a daily estimate of adult female mosquito abundance for each subregion and year in an average area *A*. Then, the estimated mosquito abundance was included in an *“epidemiological model”*, aimed at simulating the transmission dynamics of WNV in a competent bird population. Due to their abundance and competence for WNV, as proposed by Marini et al. [25], mosquitoes of *Cx. pipiens* species were considered the only vector species. Analogously, as magpies are competent for WNV [18,27] and abundant in the Lombardy region, the avian host population was considered to grow and die with rates estimated for magpies [23]. The dynamics of the disease were simulated from May to October during 2016, 2017 and 2018 according to the data provided by entomological surveillance. For both the *entomological* and *epidemiological models*, the posterior distributions of the unknown parameters were explored following a Markov chain Monte Carlo (MCMC) method following the approach adopted in Marini et al. [25]. For both models, temperature-dependent parameters were estimated using the daily mean temperature for each quadrat, which originated from ARPA Lombardia. The full model framework is shown in Fig 2.

**Fig 2.**
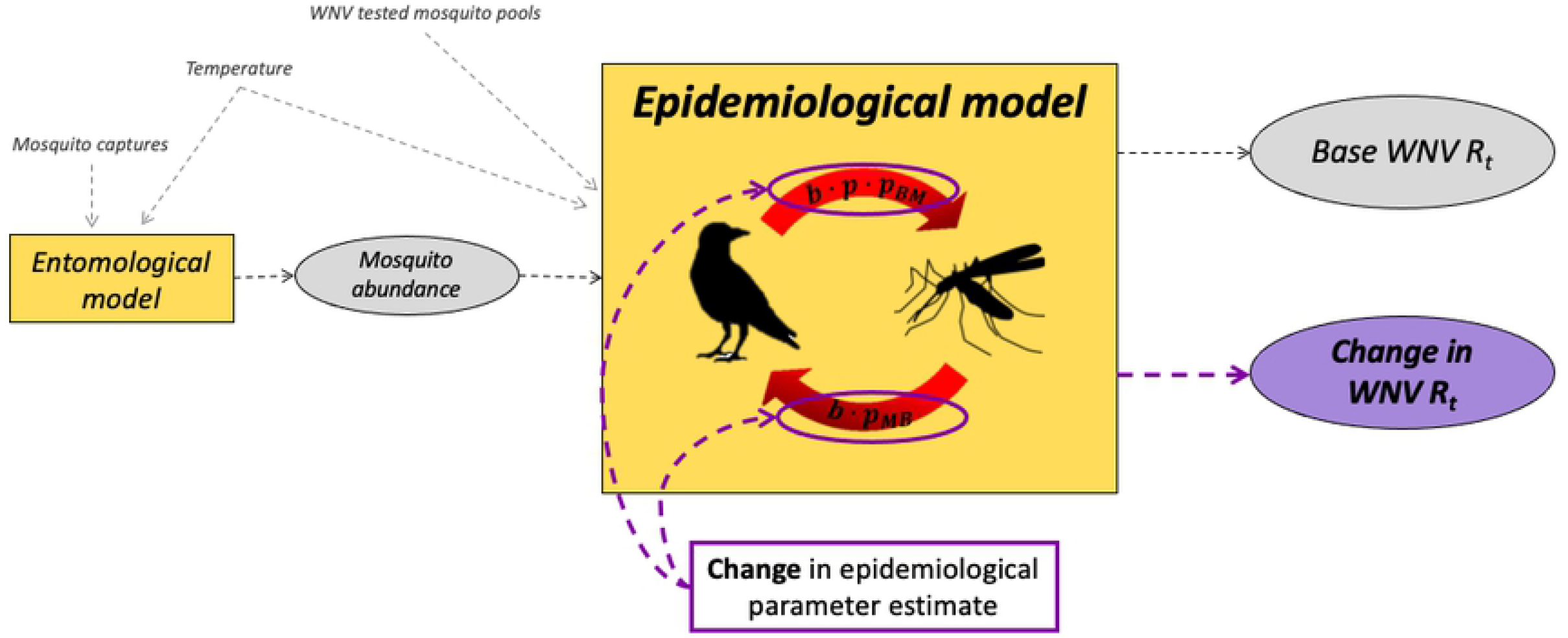
Schematic representation of the computational framework. Yellow squares: entomological and epidemiological models; grey circles: base model output; purple circle: sensitivity analysis output.

### The entomological model

To estimate *Cx. pipiens* abundance during the summer season, we calibrated the temperature-dependent entomological model presented in [25] on the recorded captures in the Lombardy region, averaged over each subregion. The posterior distributions of the unknown parameters (see below) were explored through an MCMC approach applied to the likelihood of observing the weekly number of trapped adults. Mosquito abundance was estimated for three years (2016, 2017 and 2018), starting from April up to October. The resulting mosquito abundance was then included as a known function ω(t) in the epidemiological model.

### The epidemiological model

The WNV epidemiological model is based on a system of eleven coupled differential equations, representing *Cx. pipiens* and a competent avian species divided into two age groups (juveniles and adults), and four infectious stages (susceptible, exposed, infectious and recovered). To account for the complexity of the avian population, the number of competent birds was estimated as a fraction (*a_i_*) of the total number of birds estimated to live in the area (B_0_). Birds (adults and juveniles, respectively) are considered to become infectious (*B_Ia_* and *B_Ij_*) after an incubation period and then, as a consequence of infection, recover and become immune to reinfection (*B_Ra_* and *B_Rj_*). At the start of the season (May), the avian population is considered to be fully composed of adult birds, reproducing and giving birth to juvenile birds until mid-July. Because the maturity age of birds can be considered one year [33], the newly born birds are considered juveniles throughout the entire season. Mortality due to infection in birds was neglected because of the limited mortality due to WNV infection observed in magpies in Europe [2]. Mosquitoes are considered to become infectious after a temperature-dependent incubation period (θ_M_), and they remain infectious for the rest of their life. Birds can acquire infection according to their susceptibility (*p_MB_*) and to a mosquito temperature-dependent biting rate over the bird population (*b*), depending on the proportion of mosquito bites directed to competent avian species and a function of temperature [34]. Two different biting rates were considered in the model (*b*_1_ and *b*_2_): the first for the initial part of the season (*b*_1_) and the second for the latest part of the season (*b_2_*) because of the observed shift in the mosquito biting rate between the early and late seasons [35]. The infection rate of mosquitoes was estimated by a temperature-dependent function representing the probability of WNV transmission from infectious birds to mosquitoes per bite (*p_BM_*, [23]). Viremia titres developed after infection can vary widely among bird species [16]. Avian competence is considered the probability of a mosquito becoming infectious after biting an infectious bird; thus, it was modelled as scalar (*p*) multiplying the probability of WNV transmission from birds to mosquitoes (*p_BM_*). The avian recovery rate (ν_B_) is represented by the inverse of the duration of the infectious period in birds. For both the *entomological* and *epidemiological models*, the posterior distributions of the unknown parameters (see below) were explored through an MCMC approach, applied to the binomial likelihood of observing the weekly number of positive pools given the predicted mosquito prevalence.

The complete scheme of the model and the equations describing the system are reported in the epidemiological model structure paragraph within the Supplementary Materials.

### Free parameter estimate

All biological and epidemiological parameters used for birds and mosquitoes related to magpies and *Cx. pipiens* are those reported in Marini et al. [25]. A total of 10,000 iterations of the MCMC sampling were performed, yielding 10,000 suitable sets of parameters. Simulations for WNV dynamics were then performed randomly based on the choice of 100 different sets of parameters from the estimated posterior distributions, yielding 100 different possible dynamics depending on the chosen parameter set.

The unknown parameters estimated by the MCMC method for the entomological model are:

- K_1_: density-dependent scaling factor driving the carrying capacity for the larval population at the beginning of the season
- K_2_: density-dependent scaling factor driving the carrying capacity for the larval population at the end of the season
- M_0_: the number of mosquitoes at the beginning of the season

We considered two different scaling factors for carrying capacities, as during summer *Cx. pipiens* breeding site availability might change, causing a possible increase in larval mortality, for instance, because of competition for resources with *Ae. albopictus* at the larval stage [36].

For the epidemiological model, the unknown parameters are:

- *a_i_*: the proportion of competent birds out of the total avian population in subregion *i*
- b_1_: the fraction of mosquito bites on the competent avian population at the beginning of the season (from May to mid-July).
- b_2_: the fraction of mosquito bites on the competent avian population at the end of the season (from mid-July to October).
- *i_b_:* the number of immune-competent birds in each subregion at the beginning of the first year of simulations (2016)
- *p*: avian competence, defined as the probability for a competent infectious bird to transmit the infection to a mosquito
- *p_MB_:* bird susceptibility to infection, considered as the probability for a competent bird to become infected when bitten by an infectious mosquito
- *ν_B_:* the bird recovery rate, considered as the inverse of the duration of viraemia

As a characteristic of the infection itself, all epidemiological parameters were considered constant in time and space and were estimated across all years and subregions. Only the number of immune-competent birds was considered different among years and between subregions. The number of immune-competent birds was estimated for 2016 (the first year of simulation), and for the following years, it was considered dependent on the level of avian immunity estimated at the end of the previous year. The total initial number of birds was randomly chosen in the range of 50-70 to account for the variability in avian abundance across sites and years. Mosquito WNV prevalence at the beginning of each season was randomly chosen in the range of 0-0.001 [25].

Finally, to verify whether there was a statistical relationship among unknown epidemiological parameters, we checked for their mutual correlation (Pearson correlation).

### Basic reproduction number (R_0_)

We estimated the WNV basic reproduction number (*R_0_*) following the formula proposed in [37] for vector-borne diseases. In this system, *R_0_* represents the average number of secondary infected mosquitoes over the entire transmission cycle, following the introduction of an infected mosquito into fully susceptible mosquito and bird populations. The formula takes into consideration both the number of secondary infected birds following the introduction of an infectious mosquito into a fully susceptible bird population and the number of secondary infected mosquitoes following the introduction of an infectious bird in a fully susceptible mosquito population.

The basic reproduction number is computed according to the following formula:

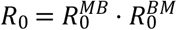

with

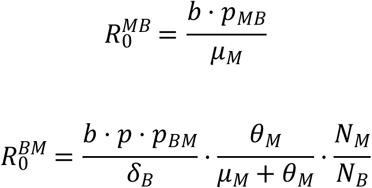

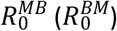 is the reproduction number of WNV and represents the number of hosts (mosquitoes) infected by an infectious mosquito (host). Following the formula proposed, the *R_0_* estimate depends on the values of epidemiological parameters as well as on the vector-host ratio 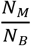, where N_M_ represents the number of mosquitoes and N_M_ the number of competent birds. Considering the wide variability of mean daily temperatures between April and October in the study area (which can easily range from 14 to 35 °C, according to ARPA Lombardia records) and the great effect that temperature has on model simulations by affecting the mosquito death rate (*μ_m_*), the probability of WNV transmission from infectious birds to mosquitoes (*p_BM_*) and the mosquito biting rate (b) [38–40], we arbitrarily chose to perform all simulations to estimate R_0_ and R_t_ by using a fixed temperature of 24 °C.

### R_t_ estimate

To estimate the potential of WNV to spread during the summer season, we adapted the formula proposed for *R_0_* to estimate the effective reproduction number *R_t_*, defined as the number of new infections caused by a single infected individual at time *t* in a partially susceptible population.

The effective reproduction number is thus computed according to the following formula:

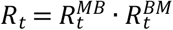

with

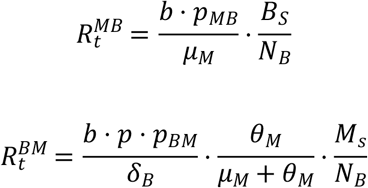

where 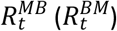 is the number of hosts (mosquitoes) infected by an infectious mosquito (host), *M_S_* represents the number of susceptible mosquitoes, *B_S_* is the number of susceptible birds, and *N_B_* is the total *M_s_* number of competent birds. As a consequence, the *R_t_* estimate depends on the vector-host ratio 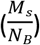 and on the proportion of susceptible hosts over the whole host population 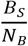.

### Transmission spread and maintenance during summer season

To investigate the transmission of WNV throughout the summer season, for each set of parameters estimated by the MCMC procedure, a daily *R_t_* was estimated from May to October, accounting for the daily mean temperature. Then, to investigate the probability of the infection being maintained and spread during the season, we estimated the monthly frequency for *R_t_* to be above 1. Hereafter, we refer to the abovementioned probability as the *spread probability*.

### Sensitivity analysis of unknown epidemiological parameters

Sensitivity analysis was performed on the following parameters:

- Mosquito biting rate (*b*)
- Competence for WNV of the bird species (*p*)
- Competent birds’ susceptibility (*p_MB_*)
- Duration of competent birds’ infectious period (recovery rate, *ν_B_*)

To investigate the effect of different parameter sets, we performed a sensitivity analysis by varying each epidemiological parameter estimate in turn and evaluating how that change affects *R_t_*. A baseline effective reproduction number 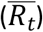 was estimated according to average calibrated parameters, and then by varying each free parameter estimate by 10%-200% (with a step of 10%), the new *R_t_* was calculated. To quantify the effect of each parameter on the effective reproduction number, we computed the ratio between the baseline 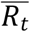 and the varied *R_t_*. We can notice that by computing this ratio, we measured the effect of changing the parameter of interest independently from the values of the others. For a deeper comprehension of the effect of a parameter variation, we also compared the monthly spread probability obtained through the base model (i.e., the number of times the estimated *R_t_* is above one divided by the total number of simulations performed every month) with the spread probability obtained with a parameter decrease of 90, 50 and 10% and an increase of 10, 50, and 100%.

### Temperature and host-vector ratio effect on R_0_

Finally, to highlight the effect of temperature on WNV spread, we performed a sensitivity analysis by varying the mean daily temperature from 10 to 30 °C (with a step of 0.1 °C) and estimating R_0_ for each temperature for each set of parameters used for simulations. Then, the WNV spread probability at different temperatures was estimated as the frequency for R_0_ to lie above 1. Although temperature affects mosquito abundance during the season, to obtain more generalizable results, we investigated the temperature effect considering only one at a time for four different values of vector-host ratios (i.e., 10, 100, 1000 and 10,000 mosquitoes/bird).

## Results

### Model calibration and fit

Parameter estimates show a very high variability for the percentage of immune birds at the beginning of the first simulated season (*i_BW_* and *i_Be_*) and for the competence (*p*), susceptibility (*p_MB_*) and recovery rate (*ν*_B_) of the bird population (see Table 1). On the other hand, the fractions of the mosquito biting rate on competent hosts (both *b_1_* and *b_2_*) and the percentage of competent birds over the whole avian population (*a_i_*) showed lower variability. A full list of the parameter estimates (and their estimated range) obtained by the MCMC approach is reported in Table 1. The 95% confidence intervals of the model predictions include 98% of the observed points, showing that despite the wide confidence intervals, the model can well describe WNV dynamics in the Lombardy region. For further details about the model fit and obtained simulations, see Fig B in Appendix S1.

**Table 1:** Estimated model parameter distributions (average and 95% confidence interval).

| Parameter | Parameter biological meaning | Estimate range<br>(2.5%-97.5% percentile) |
| --- | --- | --- |
| $B_0$ | Initial number of birds (whole avian community) | 59.95 (50.53-69.57) |
| $a_i$ | Proportion of competent birds among the whole avian community | 0.0666 (0.0016-0.2228) |
| $i_{Bw}$ | Proportion of immune birds at the beginning of the first simulated season, western subregion | 0.0445 (0.194-0.7929)<sup>1</sup> |
| $i_{Be}$ | Proportion of immune birds at the beginning of the first simulated season, eastern subregion | 0.0445 (0.1972-0.7897)<sup>1</sup> |
| $b_1$ | Proportion of mosquitoes' bites directed to the competent avian species during the early season | 0.3641 (0.1552-0.4947)<sup>1</sup> |
| $b_2$ | Proportion of mosquitoes' bites directed to the competent avian species during the late season | 0.2203 (0.0131-0.4834)<sup>1</sup> |
| $p$ | Competence of the competent avian population | 0.6191 (0.08-0.9841)<sup>1</sup> |
| $p_{MB}$ | Susceptibility of the competent avian population | 0.7752 (0.3543-0.9948)<sup>1</sup> |
| $\nu_B$ | Recovery rate | 0.3792 (0.0401-0.9253)<sup>1</sup> |

<sup>1</sup> MCMC estimate

No correlation was observed (coefficient lower than abs(0.5), P<0.05) among all unknown epidemiological parameters, except between bird susceptibility (*p_MB_*) and biting rate during the late season (*b_2_*) (corr=0.71, P<.001); bird susceptibility (*p_MB_*) and biting rate during the early season (*b_1_*) (corr=0.003, P=0.733); and bird competence (*p*) and recovery rate (*ν_B_*) (corr=0.008, P=0.416).

### Transmission maintenance during seasons

The simulated seasonal pattern of the spread probability of WNV in the mosquito population shows an initial value of the probability equal to 0.7 in May followed by an increase to 1 in June and then a continuous decrease from July (0.94) to October (less than 0.02) (Fig 3).

**Fig 3.**
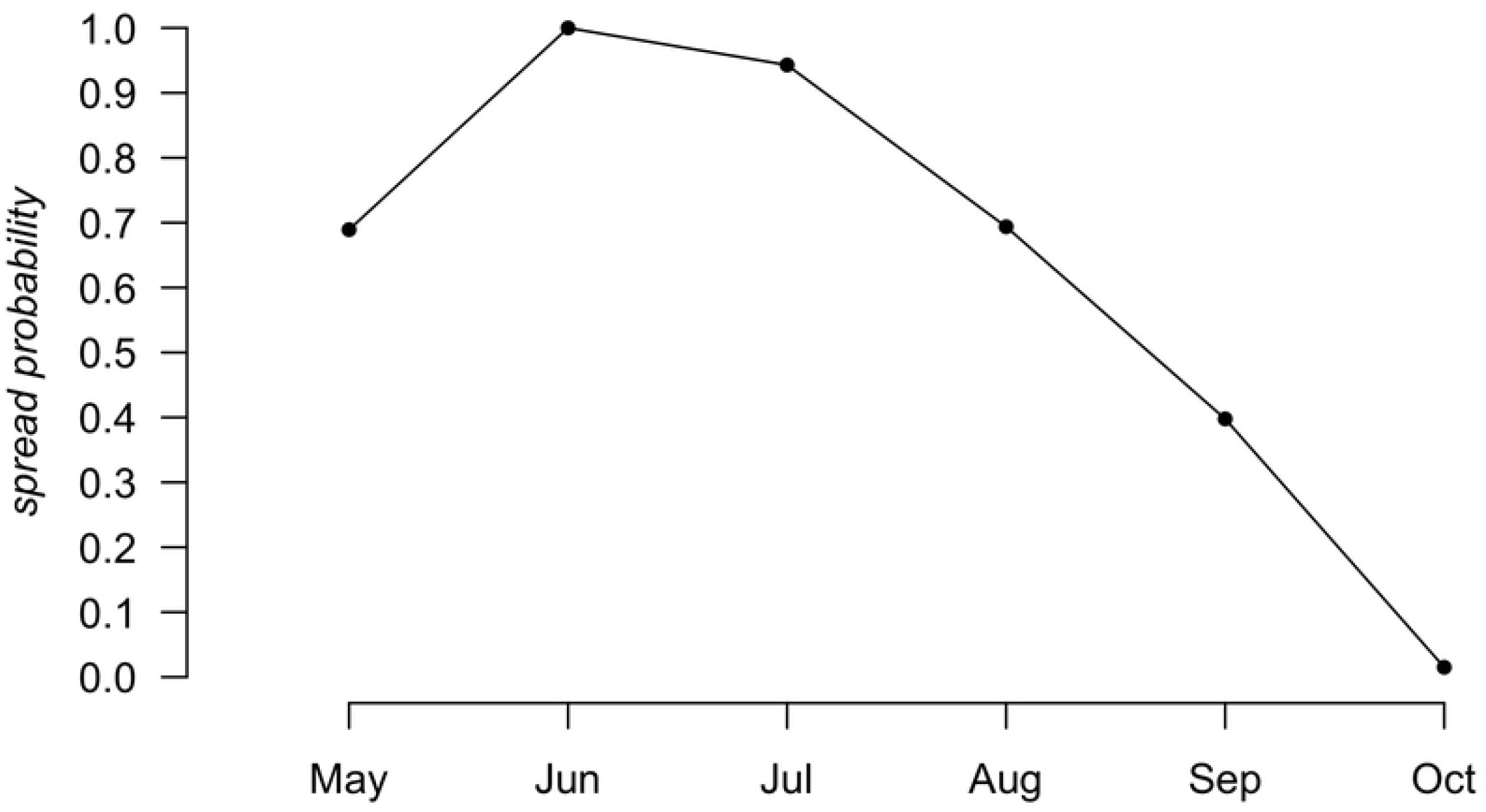
Simulated spread probability of WNV into the mosquito population during the summer season.

### Effect of unknown epidemiological parameters on R_t_

All the investigated epidemiological parameters were found to affect the relative *R_t_* (the ratio between the Rt estimated by the base model and the one estimated by changing the epidemiological parameter value in the model) estimate. Specifically, the avian recovery rate (*ν_B_*, Fig 4A) was shown to be the most impactful parameter, dramatically decreasing *R_t_* when it reached higher values (i.e., shorter infectious period); a decrease of 50% and 90% of *ν_B_* corresponded to a 2- and 10-fold increase of the *R_t_* ratio, respectively. For increasing values of *ν_B_*, the effect on the *R_t_* ratio was lower, with a decrease of 33% (50%) for an increase of *ν_B_*of 50% (100%). The mosquito biting rate (*b*, Fig 4B) is very influential as well, increasing the *R_t_* ratio with increasing values of *b*. Indeed, a 50% decrease in the *b* estimate caused a 75% decrease in *R_t_*, while a 50% increase in *b* produced a 125% increase in the *R_t_* ratio. Both recovery and biting rates had a nonlinear effect on the *R_t_* ratio, with an enhanced effect for low recovery rates and high biting rates. Bird susceptibility to infection (*p_MB_*, Fig 4C) and bird competence (*p*, Fig 4D) instead showed a smaller linear effect on the *R_t_* range.

**Fig 4.**
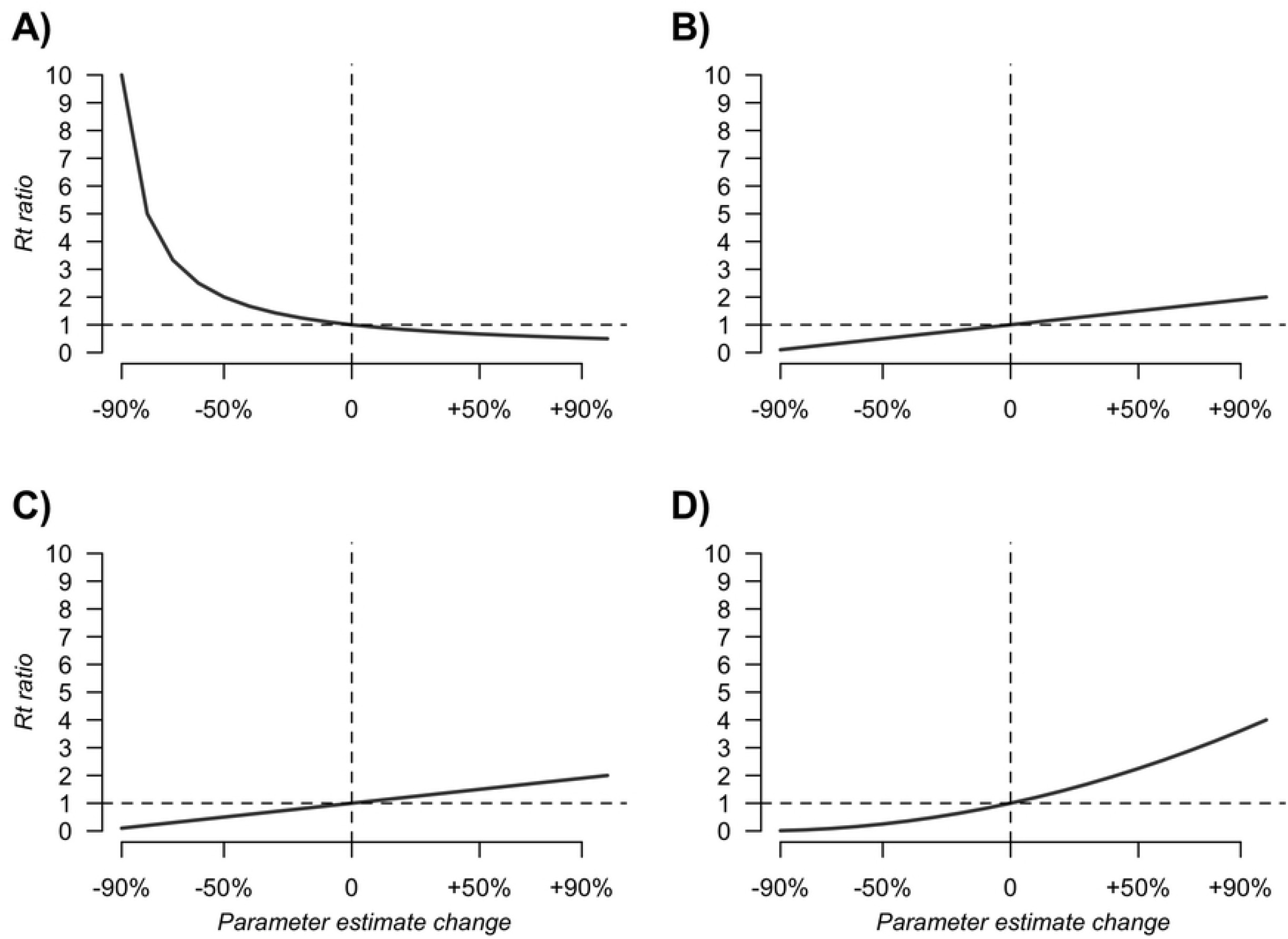
Effect of different changes in model parameters on the R_t_ ratio. Effect of A) bird recovery rate (*ν_B_*); B) mosquito biting rate (b); C) bird susceptibility to infection (p_MB_); and D) bird competence to infection (p) on the R_t_ ratio. The vertical dashed line represents the baseline parameter values (i.e., no change in parameter value), and the horizontal dashed line represents the R_t_ estimate obtained with baseline parameters.

Regarding the effect of variations in parameter estimates on the probability of spread and maintenance of WNV during the summer season (Fig 5), we observed that regardless of the variations, the month with the highest probability of WNV spread is June, followed by July, while October remains a less suitable period for WNV circulation. Specifically, the effect of parameter changes on spread probability is very low in June but is the highest in August/September. A change in mosquito biting rate (*b*, Fig 5D) highly affected spread probability, with the highest effect in July and September. On the other hand, the recovery rate (*ν_B_* Fig 5A) had less impact on the spread probability, especially for increased values of the parameter. Finally, bird competence (*p*, Fig 5B), and susceptibility (*p_MB_*, Fig 5C) showed similar results when decreased, whereas a reduction in avian competence (*p*) showed a greater effect than a reduction in avian susceptibility (*p_MB_*).

**Fig 5.**
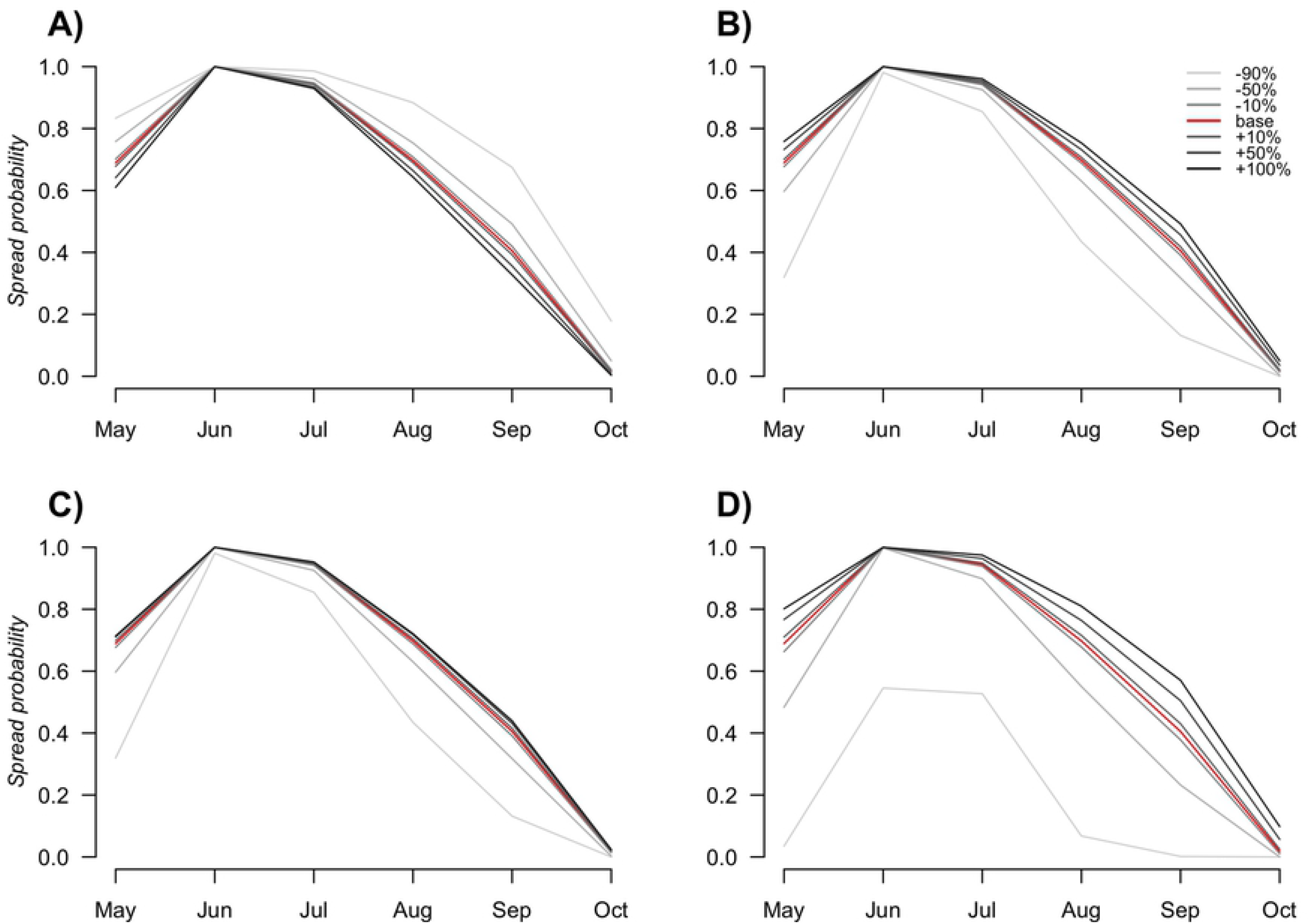
WNV spread probability in the mosquito population. WNV spread probability in the mosquito population during the summer season depended on the increase in A) recovery rate (*ν*_B_); B) bird competence (p); C) bird susceptibility (p_MB_) and D) biting rate (b) estimates. The greyscale shows from lighter to darker a change in parameter estimates of −90%, −50%, −10%, +10%, +50%, and +100%. The red line represents the baseline.

### Temperature and host-vector ratio effects on R_t_

As expected, *R_t_* is affected by temperature. Different vector-host ratios 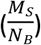 strongly impact the *R_t_ estimate*, showing a similar trend of the effect of temperature on *R_t_*, except for very low vector-host ratios 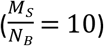. For all tested vector-host ratios, *R_t_* was always lower than that for temperatures below 14 °C, being equal to zero, in accordance with previous analyses. For 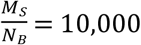, Rt rapidly increased from 0 to 1 at 14.4 °C (long dashed line). Decreasing the vector-host ratio lowered the increase in Rt with temperature, reaching 1 at 15.4 °C and 19.8 °C for 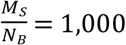 and 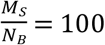, respectively (Fig 6, dashed and dotted lines). With a vector-to-host ratio equal to 10 (Fig 6, solid line), the spread probability is always below 1, reaching a maximum value of 0.43 when temperatures range between 27.1 and 28.4 °C.

**Fig 6.**
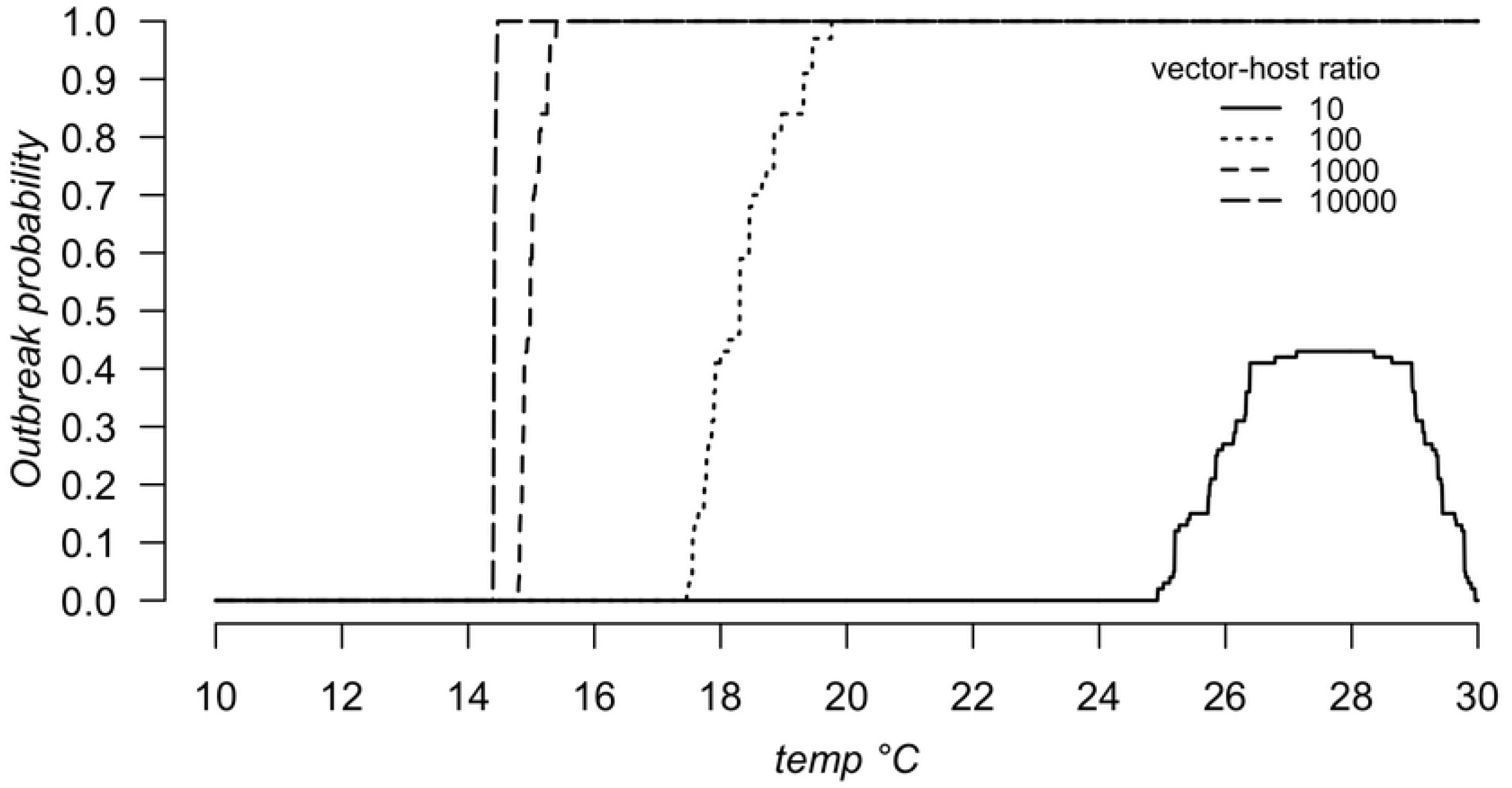
Spread probability as a function of temperature (10-30 °C) and different vector-host ratios.

## Discussion

With the present study, by calibrating a mathematical model with entomological and ornithological data gathered in northern Italy, we identified which epidemiological parameters have the largest effect on the spread of WNV. As epidemiological parameters are here used to represent transmission mechanisms, this analysis has the potential to show which mechanisms have the largest impact on infection spread, thus indicating the priorities in filling current knowledge gaps. We observed that the duration of the infectious period in birds (*ν_B_*) and the mosquito biting rate (*b*) had the largest effect on disease dynamics, whereas bird susceptibility to infection (*p_MB_*) and bird competence (*p*) showed a lower effect. In addition, according to other findings [41], temperature highly affects spread probability, not allowing WNV to spread when the daily mean temperature is lower than 14 °C. The Lombardy region, our study area, shows all suitable characteristics to allow WNV spread during summer, especially in June, when the environmental conditions seem to ensure the possibility of WNV spread in the mosquito population.

Despite the progress in health care and preventive measures, interventions to control the spread of infectious communicable diseases remain one of the main goals for public health [42,43], but this goal is often impaired by the lack of information and certainties about mechanisms driving infection transmission dynamics [17,44]. Mathematical modelling can be an efficient tool to investigate infection dynamics and transmission mechanisms (e.g., [45] and [46]), but the adoption of this approach requires robust parameter estimates to be reliable, and it has often been hampered by existing limitations due to inadequate or partial data availability. If, on the one hand, the development of field and laboratory investigations is fundamental to enhance our comprehension of spreading mechanisms, on the other hand, this process can be very long and demanding, making a prioritization of the most useful investigations essential.

Nonetheless, epidemiological mechanisms do not have the same impact on disease dynamics. Indeed, a small change in some parameter estimates can cause a large variation in model predictions, whereas large variations in other parameters can result in a smaller variation in model prediction [47]. Therefore, before designing any field study, it can be helpful to identify those parameters that have the largest impact on the infection dynamics and hence require primary attention.

WNV is now considered one of the most widespread arboviruses in the world, with human cases identified worldwide [48]. Nevertheless, there are several mechanisms that drive its spread and maintenance into wild populations, which are still unknown [17]. Many species are considered suitable as hosts or vectors, thus possibly giving different contributions to infection dynamics depending on individual and species-specific characteristics [16,18,22,49]. Moreover, the variety of involved species, the different patterns in virus transmission cycles, and the existing differences in avian community composition among different areas [12,19,50] further contribute to increasing the sources of variability in infection dynamics, thus making it difficult and expensive to collect field data required to fill knowledge gaps. Indeed, antibodies against WNV have been detected in a broad range of wild and domestic bird species worldwide, and the virus has been isolated from different avian species [12]. In addition, viraemic titres developed by different birds have been shown to strongly depend on both the host species and virus lineage [15,16,18,19]. Furthermore, mosquito feeding behaviour can change among areas and mosquito species, depending on both host abundance and mosquito feeding preference [35,51,52]. All these characteristics are suitable to drive or influence WNV spread.

In this context, the present work aimed to reveal how a change in WNV epidemiological parameters can affect our estimate and then comprehension of infection spread. First, we highlighted the importance of the bird recovery rate in driving WNV spread, showing that a change in this parameter value, especially if we consider low recovery rates (i.e., long infectious period duration), widely affects the effective reproduction number of the infection *R_t_* (i.e., the number of secondary infected mosquitoes in a given day). Changes in the bird recovery rate, despite highly affecting the estimated number of infected mosquitoes, affect the spread probability only when lowering it, supporting the importance of having long durations of infectious periods to allow for infection spread. On the one hand, these results highlight the need for a careful estimate of species-specific recovery rates to obtain a reliable estimate of WNV probability and spreading. On the other hand, they point out the need to compare species-specific rates to determine which of the investigated avian species plays the major role in spreading the infection. According to model simulations, the mosquito biting rate can also widely affect the effective reproduction number (*R_t_*) of WNV, also having an effect on the probability of the infection to spread during summer. The interaction between birds and mosquitoes is known to play a central role in disease spread [53]; moreover, it has been shown that mosquitoes can selectively choose where to feed, preferring specific species to others [35,54]. According to our results, since the effect of biting rate estimates is important and particularly high for increasing biting rate values, we can conclude that understanding which conditions and species-specific characteristics drive the probability of being bitten is critical to fully understand and predict the spread of WNV. In addition, this result again helps us to identify avian species that play an important role in WNV transmission, such as those highly bitten or preferred by mosquitoes. Furthermore, by pointing out the role of the mosquito biting rate in infection spread, this result highlights the importance of investigating the extent to which mosquitoes actively choose which species to feed on and the extent to which the biting rate is driven by species abundance. For example, in northern Italy, blackbirds are frequently bitten by mosquitoes, even if they are less abundant than other species [35]. This finding, coupled with the high number of blackbird individuals living in the area, could suggest their role in the spread of WNV. Being aware of which avian species are primarily involved in the spread of infection would help us to fill some of the current knowledge gaps and improve our understanding of WND and would allow us to efficiently estimate and predict the risk of infection for human beings and consequently to develop appropriate intervention strategies to reduce it. Avian competence, despite the high species-specific differences reported [16], seems to have a lower impact on the spread of the infection. It is assumed that to be capable of infecting mosquitoes (*Cx. pipiens*), birds need to develop viraemic titres greater than 10^5^ PFU/mL [5], but the collection of this information can be logistically demanding and hard to perform for wild birds. Moreover, experimental infections might not successfully mimic the natural infection occurring in wildlife, as mosquito inoculations of WNV in birds can result in higher viremias than needle injections [55]. Consequently, the efforts required to estimate bird competence may be greater than the benefit obtained. As anti-WNV antibodies have been detected in several bird species [15,16,31,56], a wide variety of birds may be considered susceptible to WNV infection. Bird surveillance, ongoing in several countries, can therefore be useful to help identify susceptible and potentially competent bird species. Despite that, our analysis showed that avian susceptibility to WNV infection has a small effect on both the number of secondary infected mosquitoes and the probability of spread of the disease. This finding implies that, despite being informative on the circulation of WNV in the avian population, the investigation of the WNV positivity of birds cannot be considered one of the most significant studies to be performed. Furthermore, this work supports the hypothesis that temperature and competent mosquito presence are limiting factors for WNV spread [38–41,57–59]. Indeed, according to model simulations, the R_t_ of the infection changes following a change in temperature and in the vector-host ratio. Despite that, the effect of all epidemiological parameter estimates does not change for changing temperature and vector-host ratio, making our findings generalizable and extendible to other areas with similar environmental conditions. It is also important to note that in our study area, temperatures and recorded mosquito densities always sustain WNV spread, highlighting the high human infection risk in this area. Moreover, our model shows that few initial infectious mosquitoes and a low number of birds are enough to start and maintain the infection at the beginning of the season. This finding implies that to be able to detect viral circulation early, testing of a high number of mosquitoes and birds is required. It follows that entomological surveillance, allowing faster and easier collection of numerous samples, is likely to be more informative to detect WNV early than surveillance on birds. Moreover, due to the low bird and mosquito numbers necessary to maintain the infection and considering the possible long flying distance of birds, a surveillance plan could also be beneficial in nonendemic areas with suitable climatic conditions to detect WNV introduction early in new areas.

It is necessary to note that one limitation of the present model is the assumption of having only one competent avian species. Despite this oversimplification, which could be overcome by future studies, this modelling approach has a proven ability to simulate and investigate WNV spread in nearby regions (i.e., Veneto and Emilia Romagna), suggesting its reliability despite its limitations [23,25].

In conclusion, WNV transmission and maintenance mechanisms still have some knowledge gaps, thus impairing our ability to understand and predict its spread. Among them, the duration of the avian infectious period and mosquito biting rate are the most impactful factors driving WNV circulation. Therefore, they can be considered among the most important parameters to be further studied, and their investigation could also help in determining the avian species that play the main role in WNV spread and maintenance in northern Italy. Furthermore, temperature and mosquito abundance can be limiting factors for WNV spread, and areas with suitable conditions require the design of an efficient surveillance plan to keep disease spread under control. Finally, our results, obtained through mathematical model simulations, highlight how a synergic interaction among theoretical and field research could be beneficial for a better understanding of infectious disease spread mechanisms by allowing the formulation of hypotheses to identify the most appropriate data required to cover knowledge gaps.

## Supporting information

**S1 Appendix. Supplementary information**.

**S2 Table. Recorded average entomological captures for each year and cluster**.

**S3 Table. Total number of analysed mosquito pools for each year and cluster**.

**S4 Table. Total number of WNV-positive pools for each year and cluster**.

---

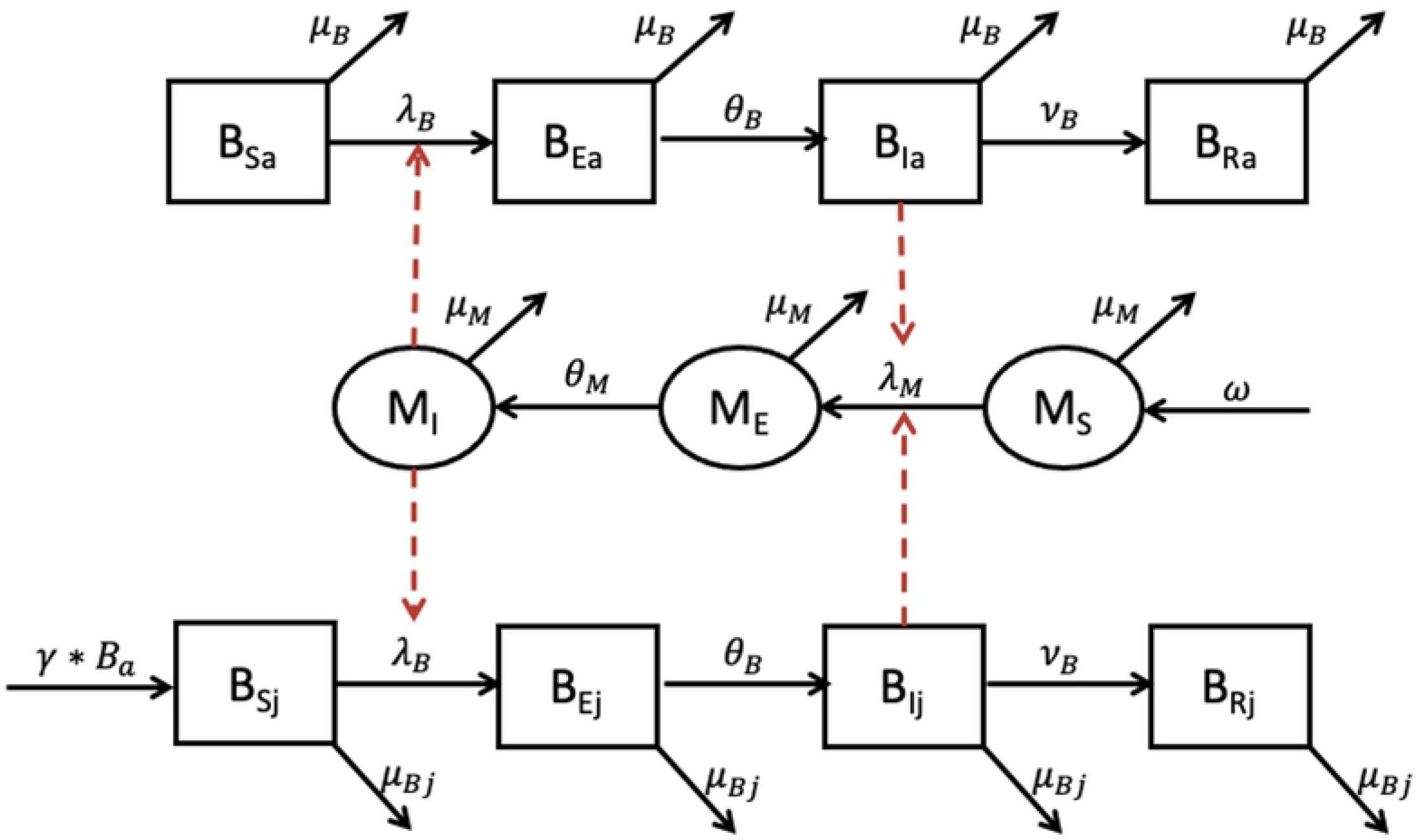
Supporting information Figure

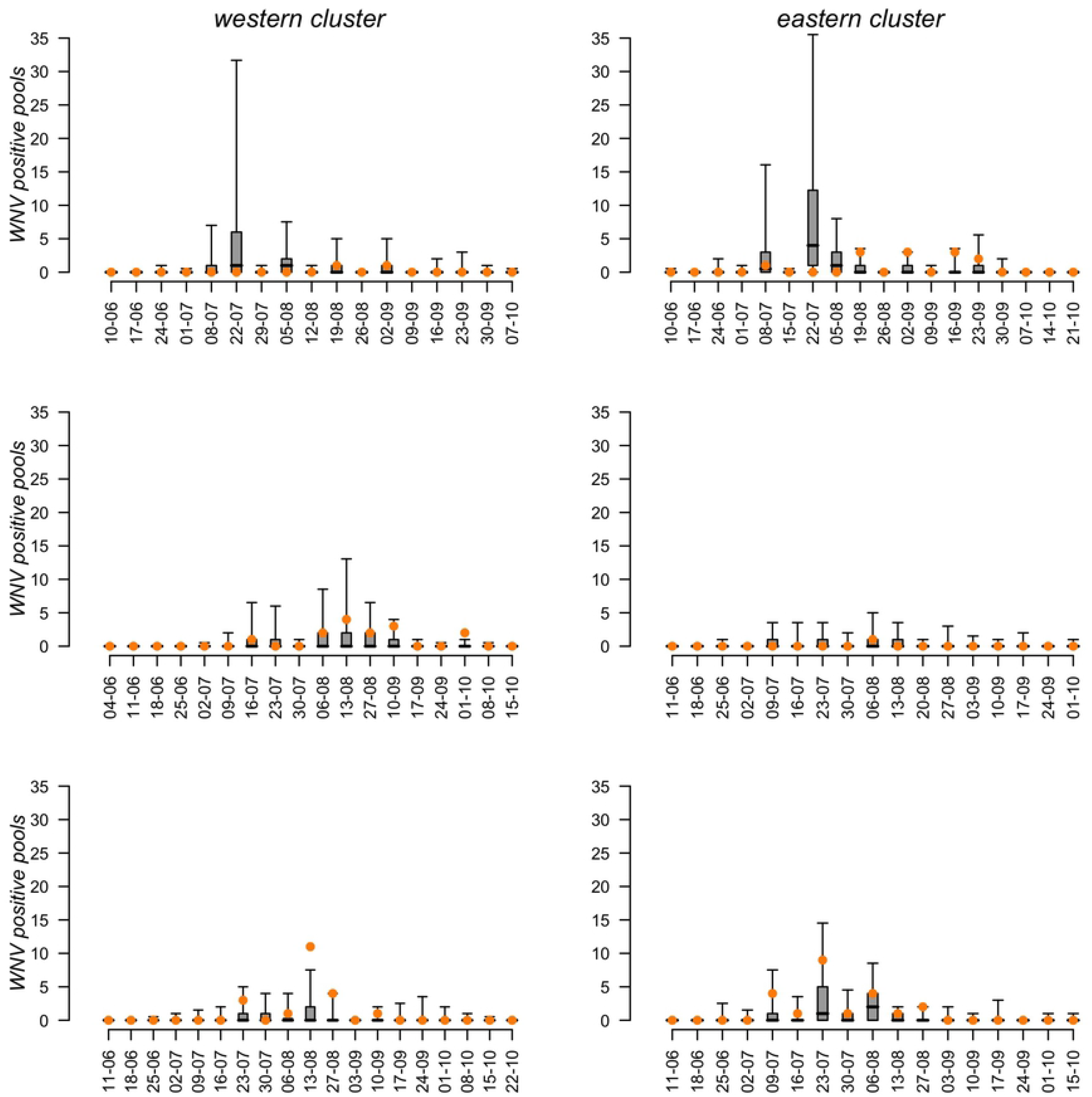
Supporting information Figure

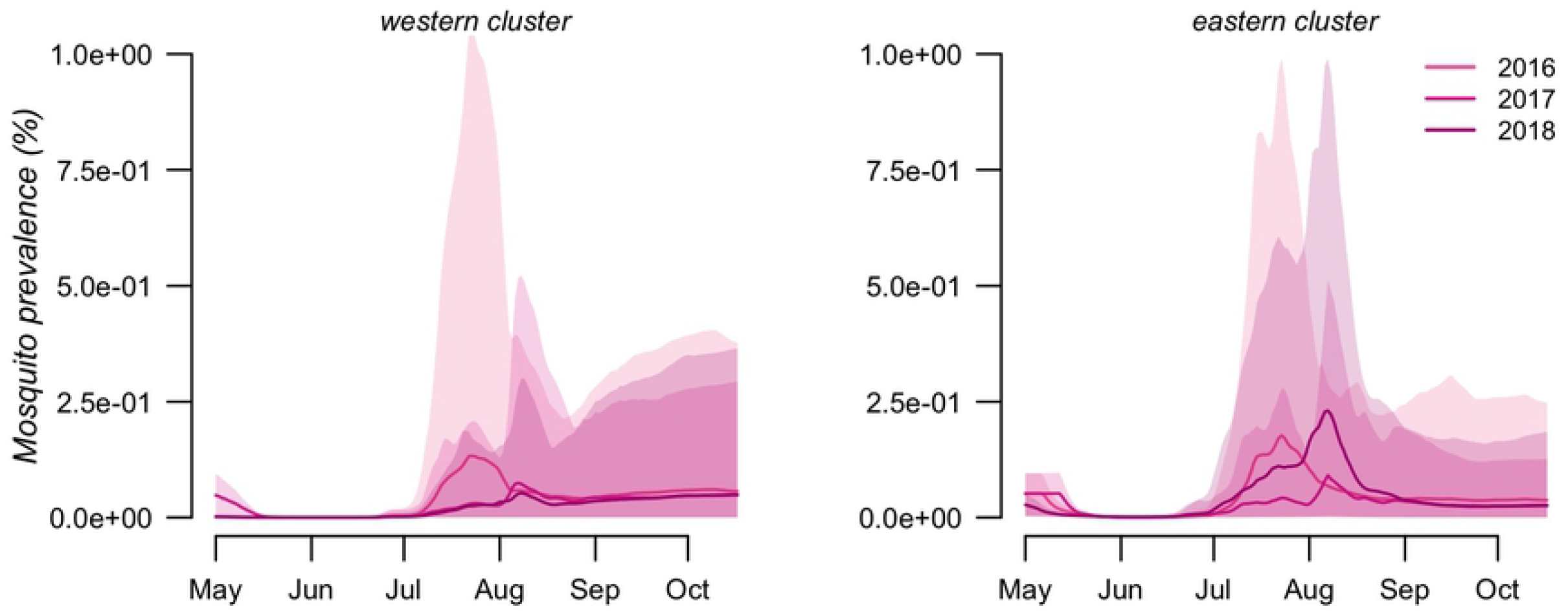
Supporting information Figure

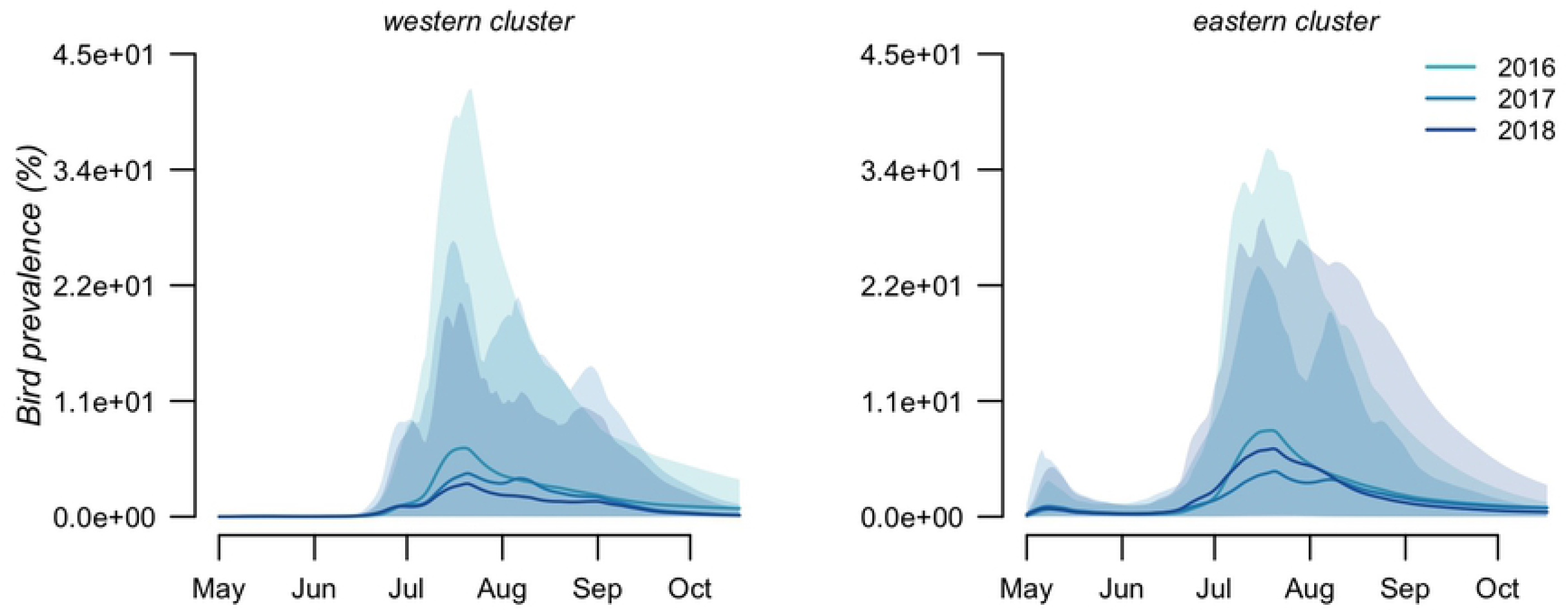
Supporting information Figure

